# Molecular Assays of Pollen Use Consistently Reflect Pollinator Visitation Patterns in a System of Flowering Plants

**DOI:** 10.1101/2020.04.14.041814

**Authors:** Aubrie R. M. James, Monica A. Geber, David P. L. Toews

**Affiliations:** Department of Ecology and Evolutionary Biology, Cornell University, Ithaca, New York, USA; Department of Biology, Pennsylvania State University, University Park, PA, USA

**Keywords:** pollination biology, bee foraging, DNA metabarcoding, relative abundance, species interactions, pollen networks

## Abstract

Determining how pollinators visit plants versus how they carry and transfer pollen is an ongoing project in pollination ecology. The differences between how pollinators visit flowers versus how they carry pollen can be appreciable, but the current tools for identifying the pollens that bees carry have different strengths and weaknesses when used for ecological inference. In this study we use three methods to better understand a system of congeneric, co-flowering plants in the genus *Clarkia* and their bee pollinators: observations of plant-pollinator contact in the field, and two different molecular methods to estimate the relative abundance of each *Clarkia* pollen in samples collected from pollinators. We use these methods to investigate if observations of plant-pollinator contact in the field correspond to the pollen bees carry; if individual bees carry *Clarkia* pollens in predictable ways, based on previous knowledge of their foraging behaviors; and how the three approaches differ for understanding plant-pollinator interactions. We find that observations of plant-pollinator contact are generally predictive of the pollens that bees carry while foraging, and network topologies using the three different methods are statistically indistinguishable from each other. Results from molecular pollen analysis also show that while bees can carry multiple species of *Clarkia* at the same time, they often carry one species of pollen. Our work contributes to the growing body of literature aimed at resolving how pollinators use floral resources. We suggest our novel relative amplicon quantification method as another tool in the developing molecular ecology and pollination biology toolbox.

## INTRODUCTION

Discovering how plant-pollinator interactions play out in natural systems is critical for understanding how such mutualisms can persist. This is because understanding resource use among mutualists is fundamental to determining the extent to which they rely on each other for population persistence (Roulston and Goodell 2011), support ecosystem functioning (Lucas et al. 2018), and mediate interspecific competition and coexistence (Johnson 2019). Two standard approaches for understanding pollination ecology in natural systems are building networks of species interactions using observations of contact between plants and pollinators, and making inferences about plant-pollinator interactions using observations of pollinator foraging behavior (Godoy et al. 2018, Manincor et al. 2020). Building networks and analyzing their topologies has been a reliable toolkit for illuminating how the community assembly of mutualistic actors operates across space and time (e.g. Alarcón et al. 2008; Bosch et al. 2009; Alarcón 2010; Tur et al. 2016; Godoy et al. 2018; Valdovinos 2019). On the other hand, studying mutualisms through the lens of individual pollinator foraging decisions can illustrate the role of pollinators in plant-plant interactions, which are the building blocks of coexistence theory and community ecology. For example, a well-documented and globally consistent trend in outcrossing flowering plants is incompatible pollen transfer between plant species, a competitive interaction that occurs when pollinators carry multiple species of pollen at once (Mitchell et al. 2009, Carvalheiro et al. 2014, Arceo-Gómez et al. 2019a,b).

Both network structure and pollinator foraging behaviors are understood using observations of pollinator visitation, and each of the following methodological approaches for documenting visitation reveals different information. First, networks of plants and pollinators can be built using observations of contact between pollinators and flowers in the field. Such observations require little work aside from identifying the pollinator and plant species at the time of observation to build a network and develop a basic understanding of how plant-pollinator communities are organized. Despite widespread application, however, evidence suggests that observations of contact between pollinators and flowers are potentially poor stand-ins for understanding how pollinators transfer pollen between plants (Mayfield 2001, Popic et al. 2013, Ballantyne et al. 2015, 2017, Barrios et al. 2016). Because of this, some studies identify and quantify pollen found on flower stigmas or pollinators using microscopy (Martin and Harvey 2017), which guarantees that plants and pollinators have interacted in a way that is relevant for both plants (pollen transfer) and pollinators (pollen resource acquisition). In some cases, however, pollens are too morphologically similar to distinguish different plant species from one another; in these situations, a third option is to molecularly interrogate pollen samples using metabarcoding (Mitchell et al. 2009, Galliot et al. 2017). Metabarcoding identifies pollen species in a sample by using single-locus PCR amplicons and high-throughput DNA sequencing to compare sample reads to a database of putative DNA sources. As a tool, metabarcoding has been applied quite successfully to identify which species of pollen pollinators carry (Wilson et al. 2010, Galimberti et al. 2014, Sickel et al. 2015, Bell et al. 2016, Bell et al. 2017, Lucas et al. 2018a,b).

Using multiple approaches to study plant-pollinator interactions has the potential to address a two-headed problem in contemporary pollination ecology: first, the desire to make sense of how plants and pollinators interact with each other, and second, the desire to compare the information we can gain using different methodological approaches in this endeavor. Field observations of plant-pollinator contact are an effective but low-resolution method to reveal species-scale interactions of plants and pollinators, and identifying pollens on individual pollinators is a higher-resolution means of understanding how plants and pollinators interact with each other with regard to pollen use and transfer. Importantly, both kinds of data can be employed to compare how field observations of contact are similar or different from observations of pollen found on pollinators; such comparisons are ultimately relevant for questions of how individuals, populations, and species mutually rely on each other as resources.

In this study, we use observations of plant-pollinator contact in the field and two different molecular methods to quantify the relative abundance of pollen sources on bee pollinators visiting a group of sympatric winter annual plants in the genus *Clarkia* (Onagraceae). This group of plants – *C. cylindrica* ssp. *clavicarpa* (Jeps.) Lewis & Lewis*, C. speciosa* ssp. *polyantha* Lewis & Lewis*, C. unguiculata* Lindl.*, C. xantiana* ssp. *xantiana* A. Gray — are sympatric in the woodland-chaparral areas of the southern foothills of the Sierra Nevada mountain range (from here, we do not use subspecies epithets). These four species of *Clarkia* rely on a handful of bee pollinators specialized on the genus *Clarkia* rather than any single species (MacSwain et al. 1973, Moeller 2005). The specialization of these bees on *Clarkia* is either incidental - *Clarkia* flower much later in the growing season than the vast majority of co-occurring flowering annual plants, and as such are the only appreciable pollen resource for bees when and where they occur – or evolutionary: some solitary bees have morphological and/or behavioral adaptations to accommodate *Clarkia’s* large pollen grains (MacSwain et al. 1973). As in many instances of closely-related groups of flowering plants, though these *Clarkia* have distinct adult phenotypes, their pollen grains are morphologically indistinguishable. Furthermore, the *Clarkia* also co-occur with each other more often than they occur alone in plant communities in their range of sympatry, and assemblages can contain one to four species of *Clarkia* (Eisen and Geber 2018).

Studies of the four *Clarkia* in their range of sympatry have alternately found signatures of *Clarkia* reproductive interference (Arceo-Gomez et al. 2016, James 2020), facilitation of pollination success (Moeller 2004), and character displacement of floral traits (Eisen and Geber 2018) in response to pollinator sharing, thereby implicating pollen transfer as an important ecological force in the system. However, these do not demonstrate how pollinators actually carry *Clarkia* pollens. Previous research has shown that the most common pollinator *Hesperapis regularis* (Melittidae) preferentially visits *C. xantiana* (James 2020). The second most common pollinator in the system, bees in the *Lasioglossum* genus (Halictidae), have been shown to either visit all *Clarkia* species at relatively the same rates (Singh 2014), or to preferentially visit *C. xantiana* and *C. cylindrica* (James 2020). Both *Hesperapis regularis* and *Lasioglossum spp.* visit *C. cylindrica, C. unguiculata*, and *C. xantiana* regularly despite their preferences, and are inconstant when foraging in mixed species experimental floral arrays, sequentially visiting multiple species while foraging (James 2020). Therefore, these pollinators may transfer incompatible pollen between plants. The third most common bee pollinator in the system, *Diadasia angusticeps* (Apidae), is more specialized on one *Clarkia* species, *C. speciosa*, and rarely visits the other species of *Clarkia* (Singh 2013, James 2020). The next step in understanding if these pollinator behaviors translate to trends in *Clarkia* reproductive interactions is to determine if pollinators collect and carry *Clarkia* pollens in the same ways that they visit flowers when foraging.

The variation in pollinator behaviors, the wealth of published knowledge of plant-pollinator interactions and natural history, the limited number of species, and the morphological similarity of pollens make this particular *Clarkia* system ideal for testing the use of multiple approaches to understand its pollination ecology. The three different approaches we use to compare *Clarkia* network topology and individual pollinator foraging decisions are: (1) field observations of plant-pollinator contacts, (2) relative read abundance (RRA) of *Clarkia* amplicons from PCR amplification, and, due to established skepticism of RRA in accurately quantifying relative abundance of species in samples, (3) a novel method we call “quantitative amplicon sequencing” or “qAMPseq”.

Quantitative amplicon sequencing is a method that quantifies the relative abundance of *Clarkia* pollen in mixed-species samples using the amplification curve of PCR as a backbone for quantification. This method identifies single nucleotide polymorphisms private to each species, and then uses PCR to amplify these regions with the goal of post-amplification sequencing, as in metabarcoding. The PCR amplification is then stopped at four different cycles in order to estimate the point at which each species’ amplification curve crosses a critical threshold (as in quantitative PCR or qPCR, Figure 1; c.f. Baksay et al. 2020). This point is used to estimate the relative abundance of each species in each sample. The goal of using qAMPseq analysis of nuclear DNA markers is to mitigate the risk of two common problems encountered when quantifying amplicons with metabarcoding: copy number bias and amplification bias. Copy number bias presents a problem for quantification because it is unclear how the abundance of a marker is related to overall pollen abundance (e.g. plastid DNA in metabarcoding using cpDNA; Oldenburg & Bendich, 2004; Golczyk et al., 2014; Bell et al. 2019). If the ratio of pollen grains to copy number is not one-to-one and/or varies among species, this could skew estimates of relative abundance in a pollen sample (Richardson et al. 2015, Bell 2019). Thus, our analysis of nuclear DNA markers means we can more reliably assume a one-to-one ratio. Amplification bias refers to the fact that the final concentration of amplicon DNA after a full PCR protocol is not necessarily directly correlated to input DNA concentration (Suzuki and Giovanni 1996; Bell et al. 2017). Instead, amplicon concentration may be a function of exhausted reaction reagents rather than the original concentrations of input DNA, especially in samples with lower initial concentrations (Bell et al. 2017, Bell et al. 2019).

**Figure 1.**
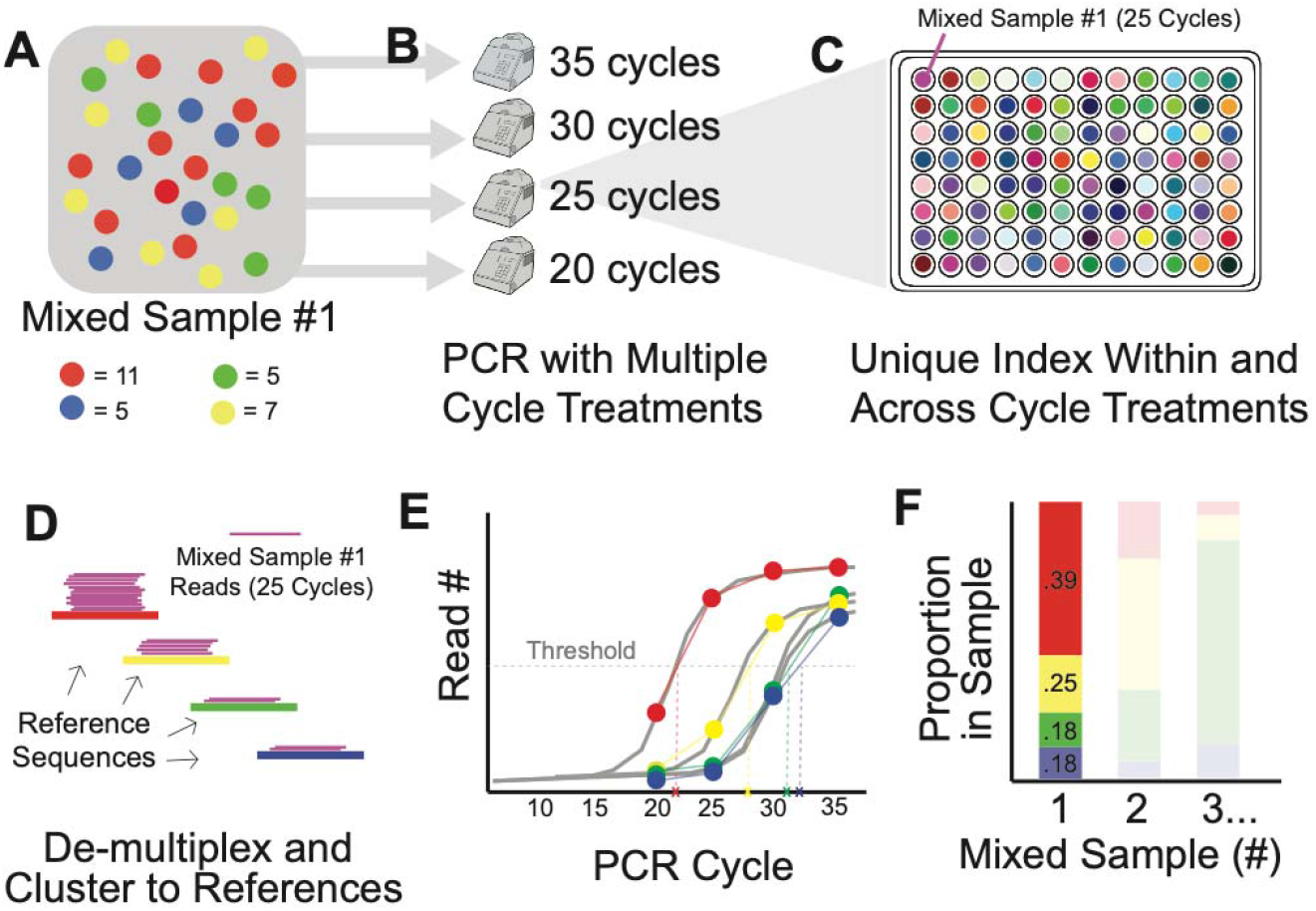
Quantitative amplicon sequencing schematic. **A** A mixed-composition sample. Each circle represents a pollen grain and species identity is indicated by color. The numbers of each pollen grain in the mixed sample are shown below (i.e. mixed sample #1 had 11 grains from red, 7 grains from yellow, etc.). **B** This mixed sample is amplified via PCR across four different thermal cyclers, each with different cycle number “treatments”. **C** Each of these samples are then uniquely indexed to keep track of sample identify and PCR cycle treatment. All samples are pooled, post-PCR, and run on a single Illumina MiSeq lane, where they are sequenced. **D** After sequencing, samples are de-multiplexed. **E** Read abundance for each species in each sample is used to calculate the Ct value from a simplified read abundance ‘curve’. **F** These Ct values are used to calculate the proportion of each species in each sample (i.e. mixed sample #1 had more red pollen grains, a lower Ct, and subsequently a higher relative proportion in the sample).

We aim to answer the following questions in this study: (1) how do network topologies of the *Clarkia* pollination system differ depending on which method we use to understand plant-pollinator interactions? (2) Do observations of plant-pollinator contact in the field correspond to the pollens that bees carry (as identified by molecular techniques)? And finally, (3) do individual bees carry *Clarkia* pollens in predictable ways, based on previous knowledge of their visitation behaviors? If each of the methods we use presents a different picture of plant-pollinator interactions, the implication would be that the choice of method is critically important in studying plant-pollinator systems. If the methods present the same or similar pictures of the system, then we will have demonstrated that results are robust to different approaches. We predict that network topologies using the three methods (observation of contact/visitation, RRA, and qAMPseq) will differ according to method: networks built using observations of contact will have fewer connections than the networks built using pollen identification and quantification from RRA or qAMPseq methods, and as a result will appear more specialized than the pollen networks. In terms of the three most common pollinator species, we hypothesize that if previously-observed trends in pollinator behavior match what they carry in their pollen samples, then *Hesperapis regularis* and *Lasioglossum spp.* will carry multiple species of pollen at once. Moreover, because these particular bee taxa have an established behavioral preference for *C. xantiana*, we expect that they will carry more *C. xantiana* than other *Clarkia* species. We also predict that *Diadasia angusticeps* will carry only *C. speciosa* pollen.

## MATERIALS AND METHODS

### Study Species and Field Sampling

The four species of *Clarkia* in this study are sympatric in the Kern River Canyon in Kern County, California. We sampled bees in 14 *Clarkia* communities throughout the four species’ range of sympatry from May-June of 2014 (Figure 2). Plant communities varied in *Clarkia* species richness (one, two, or four species of *Clarkia;* Table 1), and were far enough apart that we did not expect pollinators to meaningfully or readily move between them (Macswain et al. 1973, Zurbuchen et al. 2010). In each community, we placed four 20m long transects through patches of *Clarkia*. Each community was sampled twice on different days: once in the morning (between 8AM and 12PM) and once in the afternoon (between 1PM and 3:30PM). To sample, we walked along each transect for 20 minutes and caught bees using a sweep net when they landed on *Clarkia* flowers within one meter of the transect line (that is, the transects were 1/2 meter wide on either side of the transect line). Nets were turned inside out and quickly wiped after each bee was captured. Bees were sacrificed using ammonium carbonate. We stored, pinned, and identified bee samples to species (or in the absence of species-level resolution, to subgenus or genus) using Michener et al. (1994). We scraped the pollen contents off of all collected bees and stored each pollen sample in 90% ethanol in centrifuge tubes at −20°C. Most of the collected samples were not corbiculate bees (such as *Apis mellifera* or *Bombus sp.*), but rather bees with hairy scopae on their legs or abdomens. *Clarkia* pollen is large (100-150 μm in length) and connected by viscin threads which allows for pollen grains to be collected and deposited in sticky pollen masses made up of many pollen grains (MacSwain et al. 1973). Because of this, the vast majority of pollen samples we harvested were very loosely aggregated pollen masses that were likely more subject to bee-to-flower transfer than pollen in the tight and neatly-formed pollen balls of corbiculate bees.

**Figure 2.**
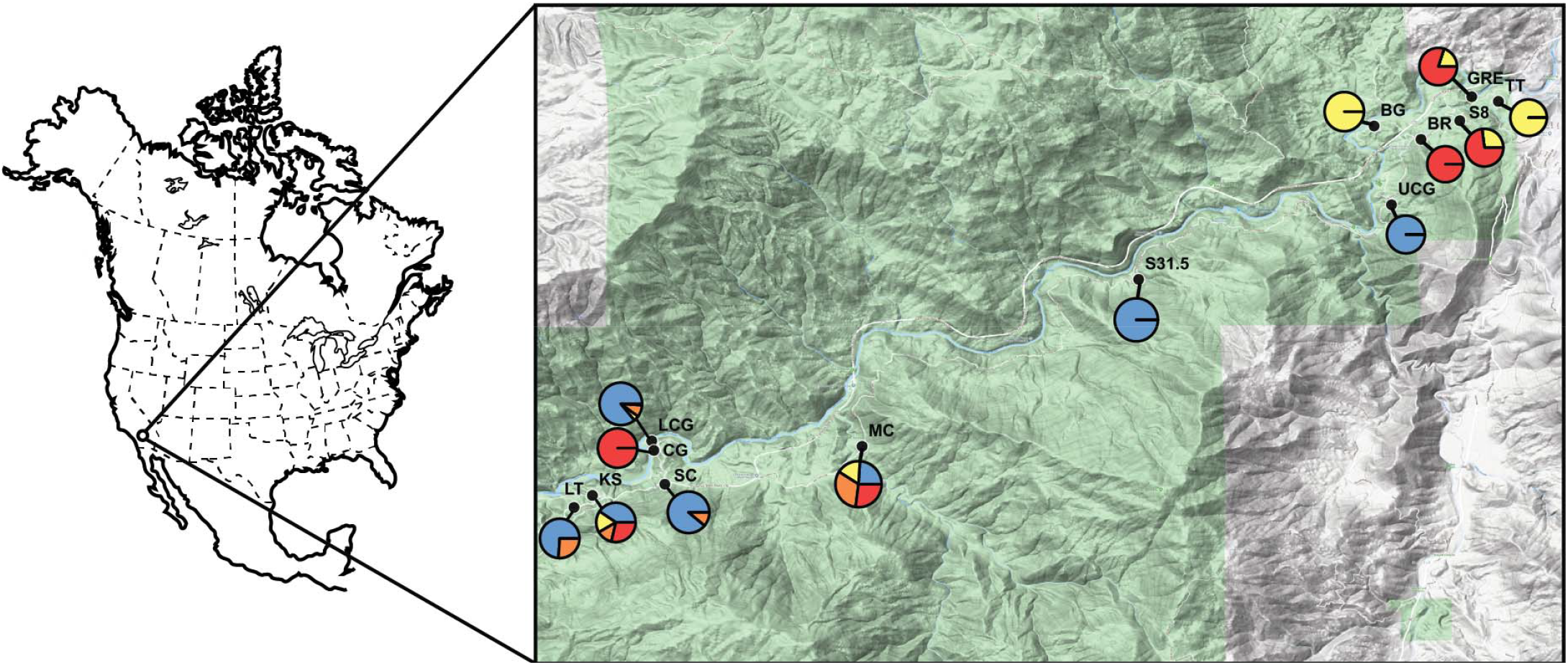
Plant and pollinator sampling locations in the Kern River Canyon in Kern County, California, USA. Pie charts show the *Clarkia* species community composition in the first round of sampling. Colors: red, *C. xantiana;* blue, *C. cylindrica;* orange, *C. unguiculata;* and yellow, *C. speciosa* (see Table 1 for details).

**Table 1.** Sites sampled in the study and the sampling schedule.

| Species composition | Site name | Dates sampled (2014) | Degree Decimal Latitude / Longitude |
| --- | --- | --- | --- |
| *C. cylindrica*<br>*C. unguiculata* | Little Tree (LT) | 15 May | 35.527197 |
|  |  | 19 May | -118.66324 |
|  | Summer Camp (SC) | 17 May | 35.531156 |
|  |  | 21 May | -118.64774 |
|  | Site 31.5 (S31.5) | 24 May | 35.566167<br>-118.56695 |
| *C. speciosa*<br>*C. xantiana* | Green Rock East (GRE) | 23 May | 35.597436 |
|  |  | 29 May | -118.51016 |
|  | Site 8 (S8) | 24 May | 35.593325 |
|  |  | 30 May | -118.51219 |
|  | Black Gulch (BG) | 29 May<br>30 May | 35.592425<br>-118.52681 |
| *C. cylindrica*<br>*C. unguiculata*<br>*C. speciosa*<br>*C. xantiana* | Lower China Gardens (LCG) | 16 May | 35.538564 |
|  |  | 26 May | -118.64998 |
|  | Kingsnake (KS) | 16 May | 35.529239 |
|  |  | 26 May | -118.6601 |
|  | Mill Creek (MC) | 17 May<br>23 May | 35.537633<br>-118.61411 |
| *C. xantiana* | Borel (BR) | 28 May | 35.590131 |
|  |  | 30 May | -118.5188 |
|  | Upper China Gardens (CG) | 18 May<br>31 May | 35.578939<br>-118.52383 |
| *C. cylindrica* | Coyote Gulch (UCG) | 24 May | 35.536933 |
|  |  | 25 May | -118.64963 |
| *C. speciosa* | Tip Top (TT) | 29 May | 35.596622 |
|  |  | 31 May | -118.50561 |

We surveyed *Clarkia* floral abundance to estimate the relative abundance of each species in the community at the end of each sampling period. To do so, we placed ½m^2^ quadrats every four meters on either side of transect lines, counting all open flowers inside the quadrats. Relative abundance of each *Clarkia* species was calculated as the number of open flowers of that species divided by the total number of flowers we counted at the survey time.

In the summers of 2015 and 2016, we also collected pollen from each species of *Clarkia* for use in testing the RRA and qAMPseq method. To do so, we collected mature anthers from all four species of *Clarkia* in various communities throughout their range of overlap. We removed pollen from the anthers and stored them in the same manner that we stored pollen samples taken from bees.

### Transcriptome sequencing from greenhouse plants

To identify nuclear genetic markers that could distinguish the four *Clarkia* species, we performed transcriptome sequencing on tissue obtained from *Clarkia* plants we grew in the greenhouse (five individuals per species; *n* = 20). We chose to use transcriptome sequencing because it was a reduced-representation genomic approach that produced long, contiguous sequences—as compared to ddRAD sequencing—that was necessary for us to develop subsequent amplicon probes. To grow plants in the greenhouse, we first cold-stratified and germinated seeds of each species of *Clarkia* in February 2016 in Ithaca, NY. Germinated seedlings were transferred into D40L conetainers (Stuewe & Sons, Tangent, 208 Oregon, USA) with a mix of 50% potting soil and 50% perlite. Once in the conetainers, seedlings were bottom-watered and grown in common conditions in a greenhouse for two months. We harvested seedlings for RNA extraction when seedlings had more than four true leaves, but before they had started flowering (March-April 2016).

Leaf tissue of seedlings was harvested and flash frozen in liquid nitrogen. To extract total RNA, we first mechanically homogenized approximately 100mg of leaf tissue with a nitrogen-chilled mortar and pestle. We then mixed the homogenized tissue with 1mL of TRIzol and 200uL chloroform, following the manufacturers guidelines for RNA isolation. We used 200uL of the isolation to a RNeasy Mini Elute silica column (Qiagen). We added 5 ul of DNAase (NEB) to the final 45 uL of the final elution, and aliquoted 20 uL of NEBNext Oligo d(t)_25_ beads to isolate mRNA from the total RNA pool. We then followed the protocol for NEBNext Ultra Directional RNA Library prep kit (NEB #E7429L). Due to low mRNA yield, we modified the protocol such that PCR enrichment included 30 cycles. We individually indexed the 20 samples and ran these on a single lane of Illumina HighSeq, using single end 100 bp sequencing chemistry.

We combined data from the five *C. speciosa* individuals to generate a reference draft transcriptome assembly using Trinity (Haas et al. 2013). We note here that many of our sequence reads derived from likely chloroplast DNA (cpDNA). Our goal was to obtain DNA markers within the nuclear genome as the presence and amount of choloroplasts within each pollen grain is unknown. We note that future studies might instead sample non-leafy tissue where possible, which will reduce the number of choloroplast reads. For the present study, we removed these reads by initially aligning the total read pools from all individuals to the chloroplast genome of a related species, *Oenothera picensis* (NCBI accession number KX118607). From the pool of reads that *did not* align to the *O. picensis* cpDNA genome, we aligned these reads from each individual to the draft *C. speciosa* transcriptome from above. We then called SNPs using the GATK pipeline, using the same set of presets as in Toews et al. (2016). We allowed for filtered out SNPs with more than 50% missing data and a minor allele frequency of less than 5%.

Ideally, our goal was to identify a single genomic region that (1) we could PCR amplify and (2) included derived, fixed SNPs for each species. To do this, we estimated Weir and Cockerham per-SNP *F*_ST_ estimates from VCFTools (Danecek et al. 2011). We generated *F*_ST_ estimates for one species compared to the other 15 individuals, and replicated this across all four species. We then determined which transcriptome contig contained multiple SNPs that had *F*_ST_ = 1 for each of the four species. We used BLAST to compare our top contig in our assembly to the nucleotide database at NCBI Genbank. The top hit was a *Clarkia unguiculata* sequence (NCBI accession number EF017402), and our contig aligned to a region that spans the *5.8S* rRNA gene, the *internal transcribed spacer 2* gene, and the *26S* rRNA gene. To amplify this region across additional samples, we used the forward primer sequence [TCGTCGGCAGCGTC]GTGCCTCGGAGATCATCTGT and reverse primer sequence [GTCTCGTGGGCTCG]GCCGTGAACCATCGAGTCTTT, with the brackets indicating the portion of the sequence (P5 and P7, respectively) that would align to our dual-indexed (i5 and i7) adaptors.

### Quantitative amplicon sequencing—An Overview

We used two methods to quantify relative input DNA from the four *Clarkia*. First, we used a common approach to quantify relative abundance of input DNA, which simply uses the relative read abundance following the full PCR. We refer to the traditional sequencing approach—using the relative read abundance of amplicons at-or-near the PCR plateau phase—as “RRA-plateau” or “RRA”. We contrast the RRA method with our method that utilizes a PCR cycle treatment, which we refer to as quantitative amplicon sequencing (qAMPseq).

The premise of qAMPseq applies the theory of quantitative PCR (qPCR, A.K.A. real-time PCR) with the ability to individually index, multiplex, and sequence hundreds of metabarcoded samples (Figure 1). Quantitative PCR analysis uses a pre-determined threshold when the PCR reaction is in an exponential phase of amplification, because the PCR cycle where a reaction product moves into the exponential phase is directly related to the starting DNA concentration, unlike the plateau stage (Kubista 2005). Quantitative PCR uses fluorescence (e.g. TaqMan chemistry) quantified throughout thermocycling to determine the ‘cycle number’ where the product fluorescence is higher than a background level, as the product is in the exponential amplification phase. The estimated number of PCR cycles when the product hits this threshold is known as threshold cycle (Ct). This Ct value can be compared across samples to compare starting DNA concentrations.

In qAMPseq, we generate the same PCR amplicon in quadruplicate, with the same starting conditions, but across different PCR cycling numbers (e.g. 20, 25, 30, and 35 cycles; Figure 2B). Subsequent cleanup and indexing steps preserve the relative DNA amounts in each of these reactions, which are then individually indexed (i.e. each original sample has four unique indexes, which correspond to the different cycle ‘treatments’) and pooled and sequenced with all other samples (Figure 2C). Samples can then be de-multiplexed (Figure 2D) and, within each sample and treatment, reads are assigned to predicted taxonomic units (“OTUs”; in this case, the four *Clarkia* species). The read abundance across each sample and OTU can be used to calculate Ct (Figure 2E), and a robust value of the relative contribution of input DNA (Figure 2F).

### Pollen DNA extraction and amplicon library preparation

Pollen DNA was extracted from sample pollen samples (2015) and anther pollen (2015 and 2016) using a CTAB-Chloroform DNA preparation protocol (as in Agrawal et al. 2013), and stored at −20°C until amplification and quantification. We first quantified the DNA concentration in each sample using a Qubit fluorometer, and diluted each DNA sample to ~2 ng/uL. We also created standard dilutions from 1:10 to 1:10000 in triplicate from a single sample of known origin. We assayed 152 unknown origin pollen sample DNA samples split between two sets. Each set included pollen DNA from 76 unknown samples, as well as the same 8 DNA samples of known origin (two from each species), and 12 samples from the standard dilution in triplicate.

Each set of 96 was then transferred to four identical 96-well plates, where we ran a PCR amplification. We conducted 10uL reaction volumes, including: 6.4 uL of ddH_2_0, 1 uL of MgCl2, 1 uL of dNTPs, 0.2 uL of each forward and reverse primers (above), 0.1 uL (0.25 units) of JumpStart Taq (Sigma-Aldrich), and 1 uL of template (at 2 ng/uL). For each set of four plates, we then used four identical thermal cyclers to run the following protocol simultaneously: 94°C for 3 minutes, and then for plates 1, 2, 3 and 4 we had 20, 25, 30, or 35 cycles of 94°C for 30 seconds, 55°C for 30 seconds, and 72°C for 1 minute, respectively. We then used a final extension time of 5 minutes.

This resulted in eight 96-well plates—four for each set—representing the different cycle treatments. We cleaned up each reaction with 1.8X volume SeraPure beads: 10uL of sample with 18uL of beads, and performed two 70% etOH washes. We eluted in 20 uL of resuspension buffer (Illumina).

We then ran an individual indexing reaction for each sample within each set (i.e. 384 randomly chosen, unique indexes for each set). The 20 uL indexing reaction included 4 uL of ddH_2_0, 10 uL of HiFi Master Mix (KAPA Biosystems), 1 uL of each the forward and reverse i5 or i7 indexes, and 4 uL of template DNA from the amplification step. This was run with the following thermal cycling conditions: 95°C for 3 minutes, 98°C for 30 seconds, followed by 8 cycles of 98°C for 30 seconds, 63°C for 30 seconds, and 72°C for 30 seconds. We had a final extension time of 3 minutes.

Within each sample set, we pooled 5uL of each indexed sample from across the four-cycle treatments, resulting in one plate for each of the two sample sets. As before, we used a 1.8X SeraPure bead cleanup for the 20uL pooled samples, and completed two 70% etOH washes. We eluted samples into 20uL of resuspension buffer. An equal volume of each sample was pooled—within each set—into the final library. We sequenced each of the two final libraries separately across two lanes of an Illumina MiSeq, with 2×150 paired end sequencing chemistry.

### Bioinformatics

Demultiplexing resulted in 1,536 individual fastq files (192 samples across four cycle treatments with forward and reverse reads). We used zgrep in bash to identify sequence motifs unique to each of the four species, combining forward and reverse read counts (Supplemental information).

We generated a standard curve by combining results from across the two sets (Figure S1). As discussed, the critical number to determine relative abundance in qPCR is the Ct value. Because qAMPseq data do not directly yield amplicon counts at the end of every cycle, we did not have direct knowledge of the exact shape of the PCR curve – the important step in determining relative abundance. To determine the cycle when samples crossed a Ct value required using a different approach: first, we log-transformed read counts associated with each of the four-cycle points for which we quantified amplicons. Log-transformation of a PCR curve theoretically results in a linear relationship of cycle and amplicon number. We took advantage of this by determining the slope of the amplification line, i.e. 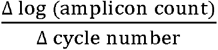. We set our Ct number as log(10,000 reads), and used it in a simple equation to determine the cycle that corresponded to Ct for each amplification curve of each species in each sample. This is similar to qPCR, where there is an arbitrary fluorescence value set above the ‘background’ fluorescence which is consistently applied across samples. In this case, the Ct number was set by identifying a region above the read ‘noise’ and below the maximal values. We henceforth call this number the cycle count, and combine this information with the standard curve to estimate relative abundance of each species in the sample.

We also set a second threshold unique to our method. If after 35 cycles a species in a sample had fewer than 10,000 reads, it was assigned a cycle count value of zero. This is because, from our known samples, we found non-specific reads can accumulate even if a species’ DNA is not present, which is to be expected in PCR to some extent. The number of non-specific reads ranged from 64 to 16,227 reads in our known samples after 35 cycles (median read count of 724). By contrast, the on-target read counts after 35 cycles from our known samples ranged from 19,770 to 744,218 reads (median read count of 135,921). Thus, the intent of this conservative threshold is to avoid falsely identifying species presence in a sample due to noise inherent in the method. We calculated cycle numbers and performed all of the following analyses in R version 3.5.2 (R Core Team, 2018).

### Comparing Network Topologies

We constructed three networks of plants and pollinators: one network with observations of the *Clarkia* species bees were caught on, and two using the *Clarkia* pollen that we identified in bees’ pollen samples (one qAMPseq network and one RRA network; we used a 5% cutoff to be considered ‘present’ in the sample for the RRA network, addressed in the following section). In the case of the plant-pollinator contact (“visitation”) network, the dataset consists of the number of times each pollinator was caught on each of the *Clarkia* species. Both pollen use datasets consist of the estimated proportions of *Clarkia* pollens in each sample, rather than a single plant-pollinator connection or the presence/absence of *Clarkia* species in a pollen sample. To build the datasets for the pollen use networks, we multiplied the proportion of each *Clarkia* species in each pollen sample by 100, and rounded to the nearest whole number.

We used two different metrics to compare the visitation, qAMPseq, and relative read abundance networks. First, we used a Procrustes analysis to compare network topology for each pair of networks (a total of three analyses) using tools in the R package vegan (version 2.5.6; Oksanen et al. 2019). In Procrustes analysis, two matrices are scaled to fit each other for maximum similarity (minimizing sum-of-squares distances), and then a statistic, m^2^, is produced to measure the goodness-of-fit between the two matrices. This statistic is bounded between 0 and 1, where m^2^= 0 means the matrices are identical. The significance of network similarity is calculated using a permutation test (Alarcón et al. 2008). Second, we measured and compared network-level specialization, H2’, in each network. We use H2’ because, like Procrustes analysis, it is robust to differences in the number of interactions (Blüthgen, Menzel & Blüthgen 2006). Values of H2’ are between 0 and 1, where higher H2’ values indicate that a network contains more specialized relationships between plants and pollinators, and lower values indicate the network has more generalized relationships. Network specialization should be the same in the visitation versus molecular networks if pollinators carry *Clarkia* pollen at the same rates that they visit *Clarkia*. Networks were built and H2’ was calculated using the package bipartite (version 2.14; Dorrman et al. 2009).

### Individual foraging and pollen loads

We used both the qAMPseq method as well as the RRA method to determine the extent to which bees carry multiple species of *Clarkia* pollen at once. In qAMPseq, we determined presence/absence of *Clarkia* in our samples by asking simply if the cycle values were nonzero (present) or zero (absent) for each species of *Clarkia*. In contrast, RRA yields relative reads after full amplification (35 cycles). To determine presence/absence of *Clarkia* with RRA output, we used three different sample proportion cutoffs to determine the presence of a species in samples: 0%, where presence was defined as any nonzero read count; and 5% and 10 % cutoffs, where presence was defined as anything above 5% or 10%, respectively. We used these three different cutoffs due to the fact that the exact percentage is arbitrary, and we wanted to account for potential differences in our RRA results due more or less conservative presence/absence read count thresholds. The 5% threshold was used as the midpoint because there is some support for using this threshold in the literature (Trevelline et al. 2018).

We compared the number of species in bee pollen samples to the amount of each flowering *Clarkia* species present when bees were collected. When we sampled bees in *Clarkia* communities, the communities contained one to four species of flowering *Clarkia*. If bees forage at random, then we expect their pollen samples to contain the same number of species as the communities in which they were captured. To test this, we tallied the number of bees caught in communities with one to four flowering *Clarkia* species, as well as the presence/absence of *Clarkia* species in each pollen sample (using our four different metrics: relative read abundance with 0%, 5% and 10% sample proportion cutoffs, (RRA, RRA5 and RRA10), and qAMPseq). We ran a Pearson’s Chi-squared test to determine if the proportion of samples containing one to four species of *Clarkia* pollen matched the proportion of bees caught in communities with one to four species of flowering *Clarkia*. If bees are inconstant when pollen foraging, these proportions would be the same, and the test would return a non-significant result.

Preference for different *Clarkia* species was estimated as the difference between the relative amount of a species’ pollen in a sample and the relative amount of that species’ floral abundance in a surveyed *Clarkia* community where the bee was captured (as in James 2020). This measure of preference can only be calculated for communities with more than one *Clarkia* species, because there is not an available ‘choice’ to make between plants in single-species communities; as such, we only calculate preference using the samples from communities with more than one *Clarkia* species. The calculation yields a value between −1 and 1 for each *Clarkia* species in each pollen sample. Positive values indicate that pollinators carry a *Clarkia* species’ pollen more frequently than it is represented in the community, values of zero indicate that bees do not preferentially forage for any species, and negative values indicate that pollinators carry a *Clarkia* species’ pollen less frequently than it is represented in the community. We calculated preference using values generated by RRA with a 5% sample proportion cutoff and qAMPseq. We used a paired t-test to determine if there was a significant difference in estimates of preference using qAMPseq versus RRA. We then ran an ANOVA and Tukey’s Honest Significant Difference test to determine if pollinator preference for *Clarkia* species were significantly different from each other, and t-tests to determine if pollinator preferences were significantly different from zero.

Finally, for each *Clarkia* species, we also summed the number of bees carrying its pollen, weighted by the proportion the *Clarkia* species was represented each pollen sample. This weighted value reveals the relative (but not absolute) amount that each *Clarkia* species was used by each pollinator.

## RESULTS

In total, we analyzed 192 pollen samples, 40 of which were samples of known pollen contents from field-collected *Clarkia* anthers, and 152 of which were pollen samples of unknown composition harvested from bees in 2014. Sequencing resulted in 45,847,334 reads, 94% of which aligned to one of the four *Clarkia* species reference sequences. All reads in known samples post-amplification were consistent with the known composition of *Clarkia* in the sample (Figure S2), barring one sample with a small number of reads. We attempted to analyze the contents of all 152 pollen samples, but two contained pollen in such low amounts they were excluded.

The final measurement of RRA with no cutoff returned all four species of *Clarkia* in 100% of the samples, indicating the unlikely result that all pollinators are not only completely random foragers, but visited all four *Clarkia* species - even when collected in communities containing fewer than four *Clarkia* species. Not only is this result unlikely based on the biology of the system, but it is exceedingly rare that any quantitative analysis with relative read abundance would use raw read count in the analysis. As such, the rest of our reported results include only information from RRA abundance reads with 5% (RRA5) and/or 10% (RRA10) cutoffs.

### Pollen use and network comparison

All three networks were topologically similar (Figure 3) and statistically indistinguishable from each other (qAMPseq vs. visitation network m^2^=0.03, p=0.001; RRA5 vs. visitation network m^2^=0.03, p=0.001; qAMPseq vs. RRA5 network m^2^=0.0006, p=0.001). Overall network specialization, H2’, was highest (more specialization) in the *Clarkia* visitation network (H2’=0.38), lowest (less specialization) in the RRA5 pollen use network (H2’=0.28), and intermediate in the qAMPseq pollen use network (H2’=0.32). Differences between networks were most apparent in the less-abundant pollinators: *Apis mellifera* (Apidae)*, Bombus sp.* (Apidae), and *Megachile sp.* (Megachilidae). Each of these species was only captured on a subset of *Clarkia*, but carried multiple species of *Clarkia*. The European honeybee, *Apis mellifera*, was only caught on *C. xantiana*, but qAMPseq analysis suggest it carried both *C. cylindrica* and *C. xantiana* pollens (on the other hand, RRA5 only identified *C. xantiana* pollen in *Apis mellifera* samples). *Bombus sp.* was only caught on *C. unguiculata* but carried both *C. unguiculata* and *C. xantiana* pollens according to qAMPseq, and carried *C. unguiculata, C. xantiana*, and *C. cylindrica* according to RRA5. Finally, *Megachile sp.* was caught on *C. cylindrica* and *C. speciosa*, but carried all four *Clarkia* pollens according to both molecular methods (Figure 3).

**Figure 3.**
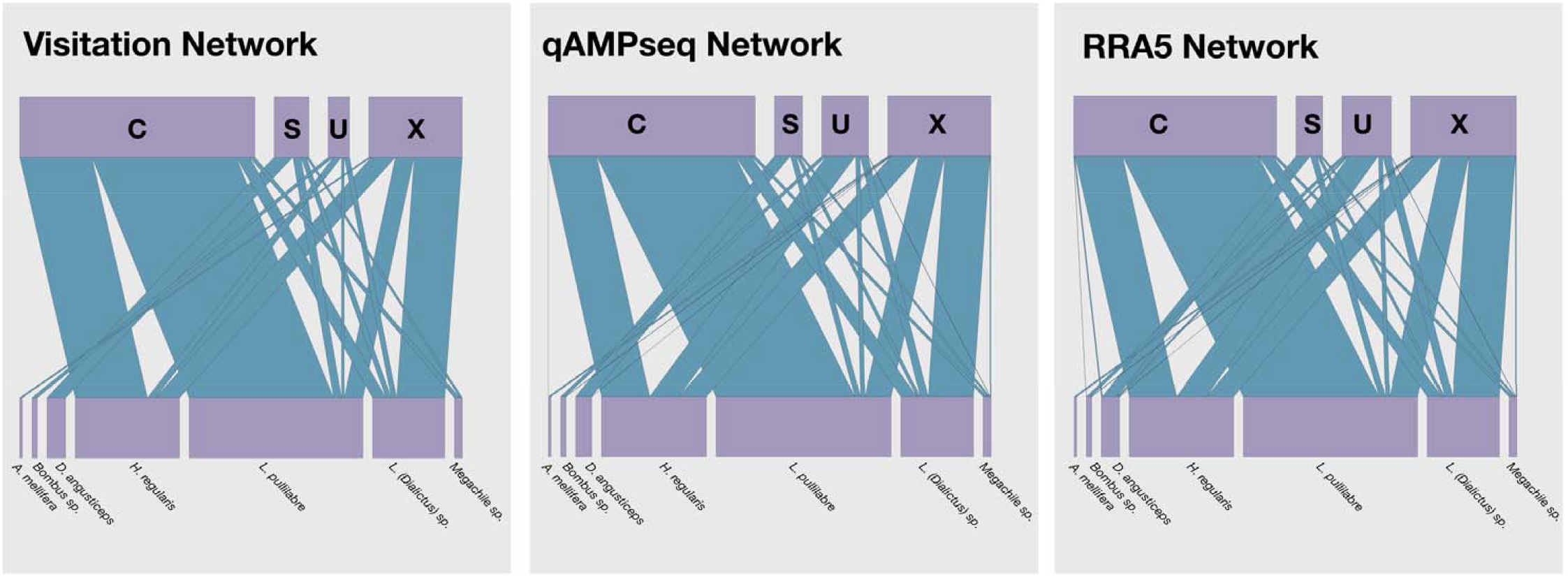
Bipartite networks of plants and pollinators. The network on the left shows the relationships of pollinator taxa (bottom row) and the plant species they were captured on (top row; C, *Clarkia cylindrica;* S, *C. speciosa;* U, *C. unguiculata;* X, *C. xantiana*). The network in the middle and on the right show the relationships of pollinator taxa and the plant species in their pollen loads using the qAMPseq method and relative read abundance (RRA5) method. The size of the bars in the top and bottom rows are proportional to the number of bees/plants in the data, and the size of the blue connecting bars are proportional to the frequency with which a plant/pollinator combination occurred.

**Figure 4.**
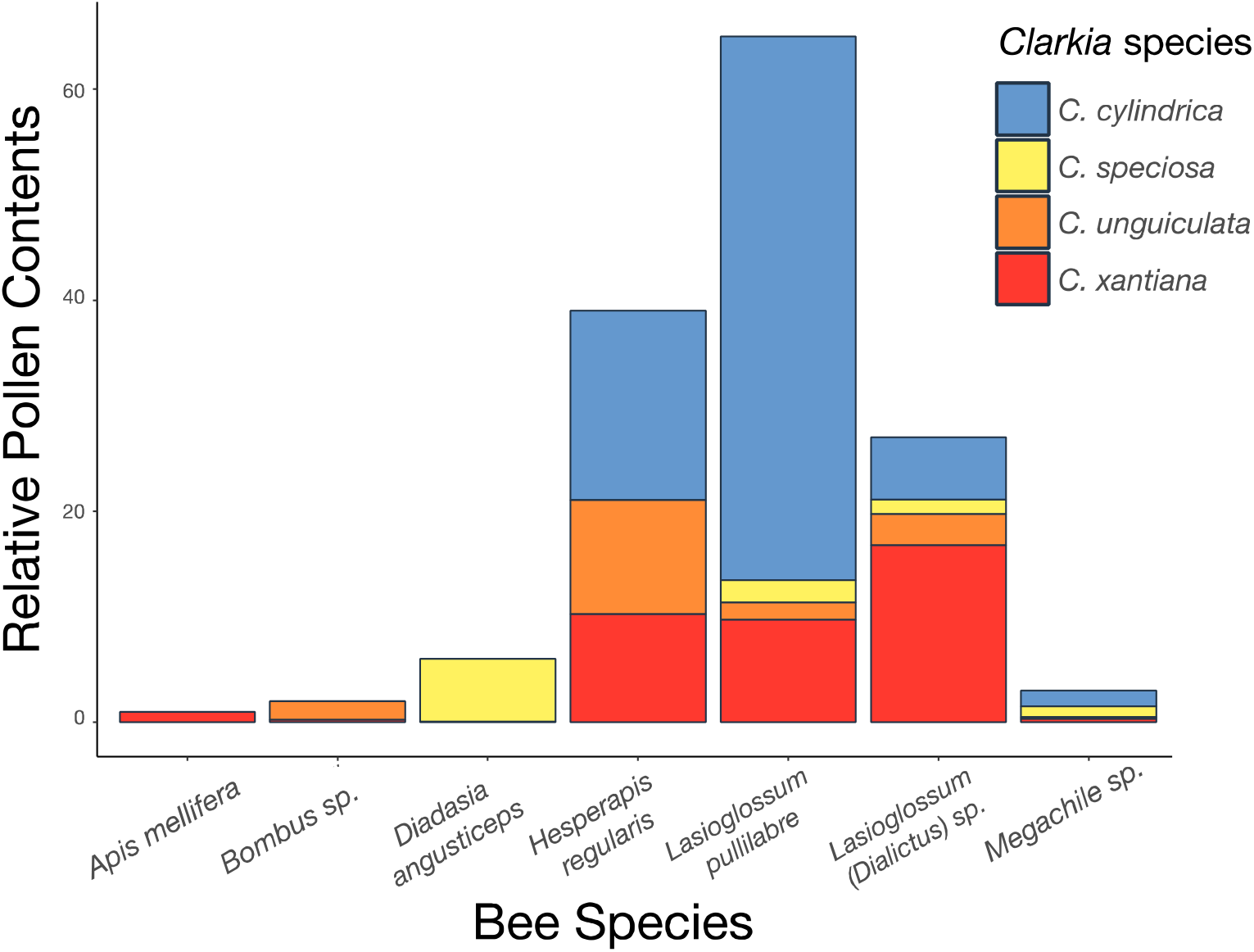
Relative abundances of *Clarkia* pollen on pollinators. *Clarkia* present in a pollen sample were weighted by their relative abundance in the sample and then summed to yield a value for that species’ relative pollen contents for each pollinator. Larger bars indicate higher pollinator counts, where *Lasioglossum pullilabre* was the most frequently caught pollinator, followed by *Hesperapis regularis, Lasioglossum (Dialictus) sp.*, and *Diadasia angusticeps*.

### Pollen sample species composition

Relative abundance measurements of *Clarkia* in pollen samples were largely the same between qAMPseq, RRA5, and RRA10, and most similar between the qAMPseq and RRA10 measurement methods (Table 2 and Figure 5; Panel B). Of the 150 pollen samples we analyzed from bees, 21 out of 150 bees (14%) were caught on *Clarkia* flowers that were different from the majority of the *Clarkia* pollen found in their pollen samples, indicating some amount of inconstant foraging (Figure 6). The contents of the pollen samples, however, frequently show that pollinators carried one species of pollen at a time. Estimates of single-species pollen samples varied among methods, with 66% (RRA5), 74% (RRA10), or 76% (qAMPseq) of samples containing only one *Clarkia* species. This indicates a striking level of pollinator fidelity, emphasized by the fact that 70% of bees we sampled were captured in multi-species *Clarkia* communities. Furthermore, the Pearson’s Chi-squared test comparing the number of bees from communities with one to four flowering *Clarkia* species versus the number of pollen samples with one to four flowering *Clarkia* species was significant (X^2^ (3)=77.05, p<0.001), confirming that even in diverse *Clarkia* communities, bees often carried only one kind of pollen (Figure 5, Panel B; Table 2).

**Table 2.**
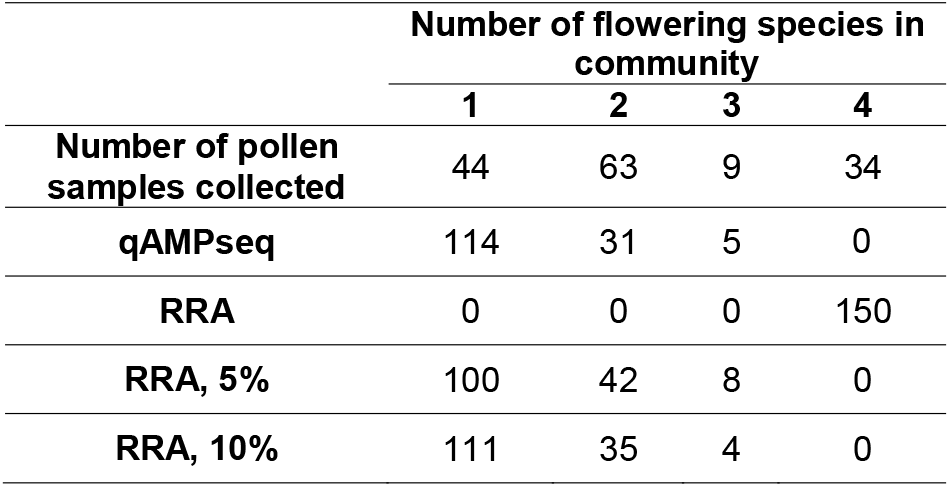
The number of pollen samples collected in communities, and the number of flowering species in the pollen samples estimated with different methods – quantitative amplicon sequencing (qAMPseq) and relative read abundance (RRA) at different sample proportion cutoffs.

**Figure 5.**
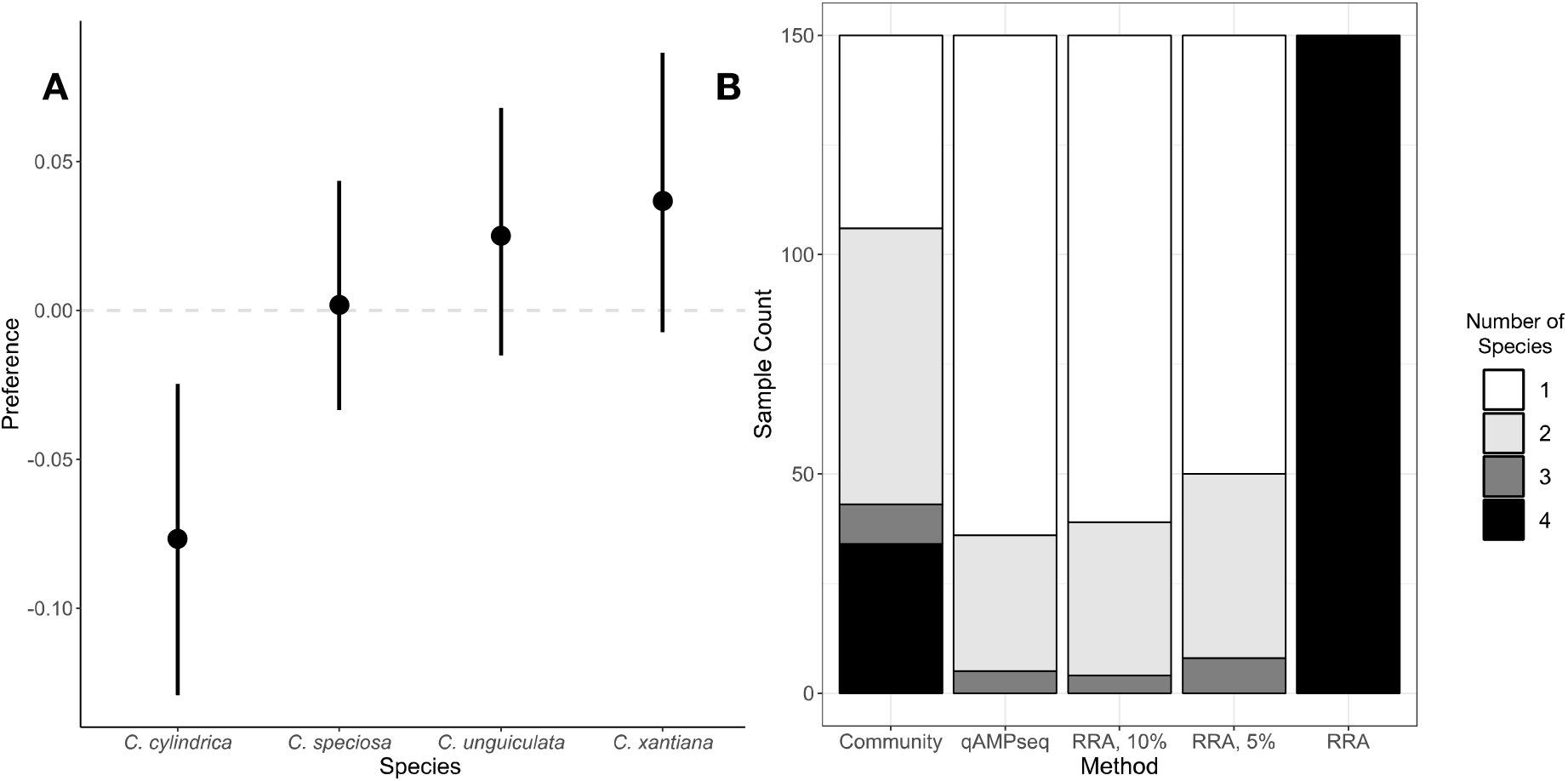
**A** Preference for different *Clarkia* pollen based on samples taken from foraging bees. Preference was calculated as the difference between frequency of a *Clarkia* species blooming in the community and frequency it was represented in bees’ pollen samples. Mean preference is plotted with a 95% confidence interval, and was estimated using quantitative amplicon sequencing. Estimates of preference did not significantly differ between methods (not shown). **B** Bar plots showing the number of *Clarkia* species in pollen samples. The far-left bar shows what we would expect if bees are random foragers: for example, the 34 bees caught in communities with four co-flowering *Clarkia* species (black section of the far-left bar) should have four *Clarkia* species in their pollen samples. We estimated the number of *Clarkia* species found on bees in four ways: quantitative amplicon sequencing (qAMPseq), and relative read abundance (RRA) at different sample proportion cutoffs. Bees carried fewer species in their pollen samples than were flowering in their corresponding *Clarkia* communities. The far-right bar is relative read abundance with no sample proportion cutoff; with this method, all samples contained all four species.

**Figure 6.**
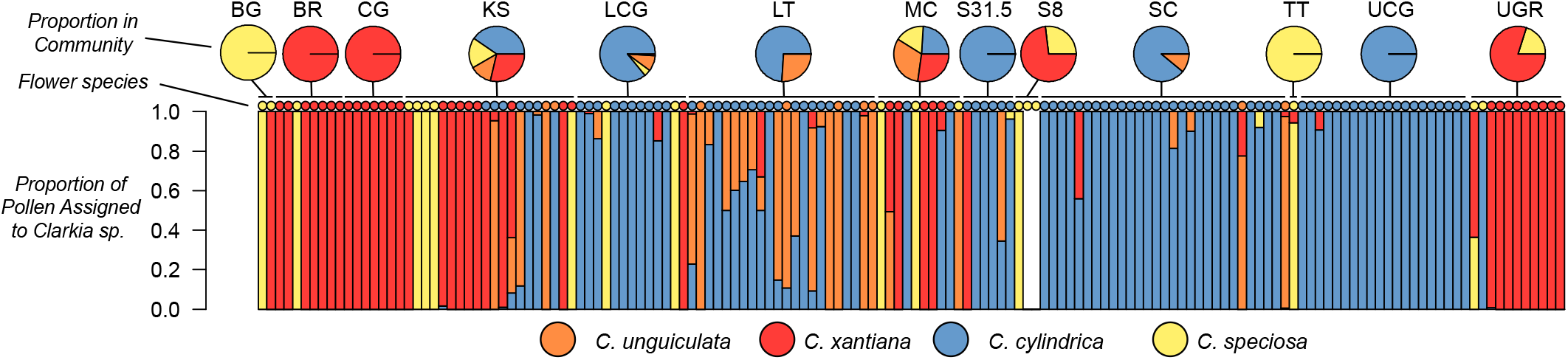
Proportion plot for each sample of pollen harvested by bees, represented by bars. Small circles above each bar indicate the species of *Clarkia* the bee was caught on, while larger pie charts indicate the *Clarkia* species composition of the community each bee came from. White bars indicate that the amount of pollen found on the bee was too low for compositional analysis.

A quarter (qAMPseq, 24%; RRA10, 26%) to a third (RRA5, 33%) of the bees carried mixed-species pollen loads, and mixtures were most often two species of *Clarkia* pollen. The most common multi-species combination in pollen samples was *C. cylindrica* and *C. unguiculata* (Figure 6). No pollen samples contained all four species of *Clarkia*. Bees carrying *C. speciosa* pollen tended to only carry *C. speciosa* pollen: when present, *C. speciosa* was the only species of pollen in the sample in 12/14 cases. Bees carrying the other three species of *Clarkia* often carried mixtures of the three (Figure 6).

Pollinator species also carried markedly different proportions of each *Clarkia* pollen. We were able to distinguish two different *Lasioglossum* taxa in our study, and found that the two identifiable taxa of *Lasioglossum* exhibited different rates of carrying each species of *Clarkia*. A putative *Clarkia* specialist, *Lasioglossum pullilabre*, carried all four species but was most associated with *C. cylindrica*, whereas *L. (Dialictus) sp.*, a likely generalist, carried *C. xantiana* at higher rates (Figure 4). Furthermore, the *Clarkia* specialist *Hesperapis regularis* carried *C. cylindrica, C. unguiculata*, and *C. xantiana* at almost equivalent rates, while *Diadasia angusticeps* used *C. speciosa* almost to the exclusion of all other *Clarkia* (Figure 4).

### Preference

The two methods we used to determine pollinator preference, RRA5 and qAMPseq, did not differ in their estimates of preference (t(607)=0.828, p=0.41). Preferences for *Clarkia* species were significantly different from each other (F(4, 604)=4.899, p=0.002). On the whole, bees carried *C. cylindrica* pollen significantly less than it was represented in *Clarkia* communities (preference = −0.07±0.0 95% CI). Bees carried the other three species at roughly the same frequency they occurred in sampled communities (*C. speciosa* t(604)=0.077, p=0.93; *C. unguiculata* t(604)=1.086, p=0.27; *C. xantiana* t(604)=1.59, p=0.11; Figure 5, Panel A).

## DISCUSSION

In community ecology, mapping the relationships between plants and pollinators is critical to understanding how pollinators contribute to plant community functioning and vice-versa. Though observations of pollinator visitation can be used to understand plant-pollinator community assembly, visitation does not necessarily correspond to pollen transfer or pollen use (Mayfield 2001, Popic et al. 2013, Ballantyne et al. 2015, 2017, Barrios et al. 2016). Metabarcoding has been critical for understanding if and when bees use certain pollen resources in plant communities (Galliot et al. 2017, Lucas et al. 2018*b*), but some studies suggest that relative read abundance from PCR could be unreliable for use in measuring abundance in mixed-species pollen samples (Bell et al. 2016, Bell et al. 2019).

In this study, we have used visitation observations and molecular methods to uncover how plants and pollinators interact. We asked if different methods reveal differences in how bee pollinators visit plants versus how they carry and transport pollen, as well as if pollinators carried pollen in predictable ways given what is known about their foraging behaviors. We discovered that the way pollinators visit plants in the field and the pollen they carry in their pollen loads are almost identical. This result is particularly striking because whereas our visitation network documents contacts between single pollinators with single flowers, our pollen networks document potentially tens or hundreds of contacts between a single pollinator and many flowers in a community. That the visitation and pollen networks correspond so closely not only in presence/absence but also in frequency of contact indicates high pollinator fidelity. This is confirmed with further interpretation of the data- our analyses show that pollinators predominantly carry a single species of pollen at a time. By quantifying relative abundance of *Clarkia* in pollen loads, we shed light on how pollinator species actually collect pollen from different *Clarkia* species: when pollinators do carry more than one species of pollen at a time, it is not at equal frequencies, as presence/absence results alone (as in standard metabarcoding analysis) might suggest.

### Pollen and Pollinators

With this study, we have contributed to the growing body of literature that uses molecular methods to understand pollen collection and transport by bees (Mitchell et al 2009; Galimberti 2014; Bell et al. 2017, Galliot et al 2017, Lucas et al. 2018, Bell et al. 2019), and in the process provided some evidence for the potential of heterospecific pollen transfer by pollinators in flowering plant communities (Mitchell et al. 2009, Arceo-Gomez et al. 2019). That said, pollen samples in our study typically comprised one species of *Clarkia*, which indicates that heterospecific pollen transfer may not be as common in the *Clarkia* system as in other flowering plant systems (i.e. Arceo-Gómez et al. 2019). The bees that did carry heterospecific pollen carried combinations of *C. cylindrica, C. unguiculata* and *C. xantiana*, suggesting that these three species are more likely to receive heterospecific pollen than their counterpart, *C. speciosa*. To decisively conclude the potential for heterospecific pollen transfer, however, more research is needed to determine how pollen is transferred between bees and flowers after pollen has been collected, as pollen contents found on pollinators does not necessarily correspond to pollen transfer.

There were two important results revealed by our pollen analysis that would not have been available using presence/absence metabarcoding or visitation observations alone. First, *C. cylindrica* was carried with higher total representation in pollen samples than other species of *Clarkia*, but it was still carried at lower frequencies than it was represented in *Clarkia* communities (Figures 4 and 5). This is probably because *C. cylindrica* has the highest average floral abundance of all the species throughout the communities we sampled (James *unpublished data*). Consequently, it is possible that pollinators might limit the competitive dominance of *C. cylindrica* via introducing pollen limitation to seed production. Second, we were able to resolve differences in foraging between *Lasioglossum* taxa. Because we collected and sacrificed the bee specimens in this study, we could identify *Lasioglossum* with higher resolution than observing visitation without sampling, and show that the taxa exhibit differences in their relationships to *Clarkia:* the putative specialist on the *Clarkia* genus, *Lasioglossum pullilabre* (Moeller 2004; Eckhart et al. 2006), carries *C. cylindrica* with higher frequency, whereas the likely generalist, *L. (Dialictus) sp.*, carries *C. xantiana* with higher frequency (Figure 4).

With this study, we were also able to better delineate *Clarkia* use by rare pollinators in our dataset. The rarest pollinators, *Apis mellifera, Bombus sp.*, and *Megachile sp.* (rare to the dataset, but not rare in the ecosystem; Moeller 2005, Eckhart 2006, and Singh 2013) all carried more species of *Clarkia* pollen than they had been observed visiting. This explains the differences in specialization of the pollinator *Clarkia* visitation network versus that of the pollen-use networks: pollinators carried more diverse pollens (and the network was therefore less specialized) than the plants we observed pollinators had visited. Given that sampling effort is a perennial issue in network analyses, pollen networks like this one and others (for example: Alarcón 2010, Galliot et al. 2017, Lucas et al. 2018) are a means to understand plant-pollinator relationships when sampling effort is constrained. We say this with caution, however: pollen analysis data should complement, not supplant, well-designed sampling methods. For example, it is highly likely that *Apis mellifera* and *Bombus sp.* use all four *Clarkia* pollens (Singh 2013), but our sample size of a few bees per species makes that impossible to say with certainty.

Other patterns in pollen use were noticeably similar to what we predicted based on standing knowledge of the pollination ecology of the system. The two *Clarkia* species most often found in multi-species pollen samples, *C. cylindrica* and *C. unguiculata*, have been shown to occur together with higher frequency than any other *Clarkia* species pair in this system, overlap in flowering time more than they do with other *Clarkia*, and exhibit pollinator-mediated character displacement in floral traits (Eisen and Geber 2018). Given the frequency with which they occur together in pollen samples, it is possible that character displacement in the floral traits of these two species could be driven by the competitive effects of heterospecific pollen transfer. Furthermore, *Diadasia angusticeps* bees carried pollen samples of single-species composition (*C. speciosa*), which corresponds to previous observations that the species is specialized on *C. speciosa* (Singh 2013, James 2020).

### Molecular methods for quantifying relative abundance in pollen samples

This study presents a novel method of meta-DNA relative abundance analysis, qAMPseq, with the motivation of avoiding some of the biases and problems that can arise when using plastid makers and RRA. However, we unexpectedly found that the two analytical approaches to quantifying pollen relative abundances in the molecular data yielded nearly identical results. The only appreciable difference between the RRA network and the qAMPseq network that we observed is that the network built using qAMPseq exhibited fewer connections overall, and thus qAMPseq was slightly more conservative than the RRA approach (Figure 3). These slight differences do not change our community-scale interpretation of the networks. Further, whereas we expected that the RRA method would yield more spurious associations between plants and pollinators, we found in almost all analyses that the methods yielded the same results. It is likely that this similarity was due in part to the fact that all of our samples were diluted to the same starting concentration of DNA (2 ng/uL). Therefore, in studies wishing to use RRA, it should be noted that RRA may not yield accurate estimates of relative abundance if DNA concentrations among samples are highly variable (Bell et al. 2017, Bell et al. 2019).

Furthermore, an important difference between our study and studies that use universal primers to amplify ITS2 and rcbL is that we had specific target species in our samples. We designed primers to amplify regions where we knew there were SNPs that differentiated our target species from each other, rather than relying on variation in ITS2 and rcbL to distinguish *Clarkia* species from each other. In the end, however, the region that ended up being the most informative among our four species included ITS2, suggesting it is an ideal starting point for future studies. It is also important to note that, because we were using closely related species within the *Clarkia* genus, our application of the metabarcoding approach may not have been subject to many of quantitative biases identified by Bell et al. (2019). These biases include copy number variation of the amplified gene, differences in DNA isolation efficacy among samples, and variation in primer amplification efficiency. If others are to use this method with primers that target a broader range of possibly more divergent species, these additional biases need to be carefully considered in experimental design.

Given the dramatic decline in sequencing costs, the sensitivity of current sequencing methods to detecting SNPs, and the potential biases that may arise when quantifying PCR products from metabarcoding in mixed-species samples (Bell et al. 2019), the qAMPseq method represents a lower-cost alternative to florescence-based qPCR (as well as being more viable; see supplement S3), and offers an alternative to relative read abundance analyses. That said, the qAMPseq protocol is time-intensive, requires that researchers identify private SNPs for constituent species in pollen samples, and could be streamlined. One example of streamlining is that it may be acceptable to perform fewer bead cleaning steps to reduce cost and time at the bench. Our protocol also necessitated four concurrently running thermocyclers, but we can envision a modification of thermocycler heating blocks that might allow for variation in the number of reaction cycles, which would allow for reactions to be run on the same machine (e.g. Schicke and Hofmann, 2007). There are also standing issues inherent with both molecular methods that we could not address: for example, our results using qAMPseq versus RRA depended on the tolerance with which we filtered raw RRA values and the critical threshold value in qAMPseq, both of which were arbitrarily selected. Furthermore, our analysis does not incorporate information about the size of the sampled pollen sample, so we cannot draw conclusions about the amount of pollen that different pollinators transport in their pollen loads. Irrespective of approach, we highlight that care must be taken in determining the concentration of pre-amplification samples with any particular primer/target combination, as well the interpretation of the resulting data.

### Conclusions

As the theories of plant species interactions continue to incorporate mutualistic, complex community dynamics (Morales-Castilla et al. 2015, Vázquez et al. 2015, Valdovinos 2019), molecular approaches can facilitate a more robust understanding of species interactions between mutualists. Here we have presented evidence that field observations of plant-pollinator contact correspond to molecular observations of how pollinators carry pollen, indicating a high level of pollinator fidelity. The networks we built using molecular approaches also show that pollinators sometimes carry mixed-species pollen balls that might lead to incompatible pollen transfer between plants. Finally, the two different molecular approaches we used yielded similar results indicating that both are valid ways of molecularly interrogating pollen samples of mixed species composition.

## Supporting information

sequence_motifs

## Acknowledgements

The authors acknowledge Steve Bogdanowicz, the Cornell Evolutionary Genetics Core Facility, the Cornell Lab of Ornithology, Bronwyn Butcher, the Lovette Lab, and the Agrawal Lab for help in method development and for providing lab space and time. Dave Moeller and Indrani Singh helped with bee identification and provided helpful natural history knowledge, and Amy Hastings helped in DNA extraction methods. We thank Julia Brokaw, who was instrumental in the field collections of bees and pollen, as well as Alyssa Anderson, who aided in data preparation. This work was supported by the National Science Foundation Doctoral Dissertation Improvement Grant program grant number 1701675 awarded to ARMJ and MAG, and NSF DEB-1754299 awarded to MAG. Finally, the authors acknowledge that the samples in this research were collected on traditionally Tübatulabal land. The Tübatulabal tribe are currently seeking legal recognition from the federal government.

## Data Accessibility

Pollen sample composition data will be archived in the Dryad digital repository.

## Author Contributions

ARMJ, MAG, and DPLT designed the study. ARMJ and DPLT performed research, analyzed data, and wrote the paper. All authors contributed to editing and revising the manuscript.

