## supplementary_materials for "Molecular Assays of Pollen Use Consistently Reflect Pollinator Visitation Patterns in a System of Flowering Plants"

**Table of Contents:**

| **Figure S1** | Page 1 |
| --- | --- |
| **Figure S2** | Page 2 |
| **Supplement S3** | Page 3 |

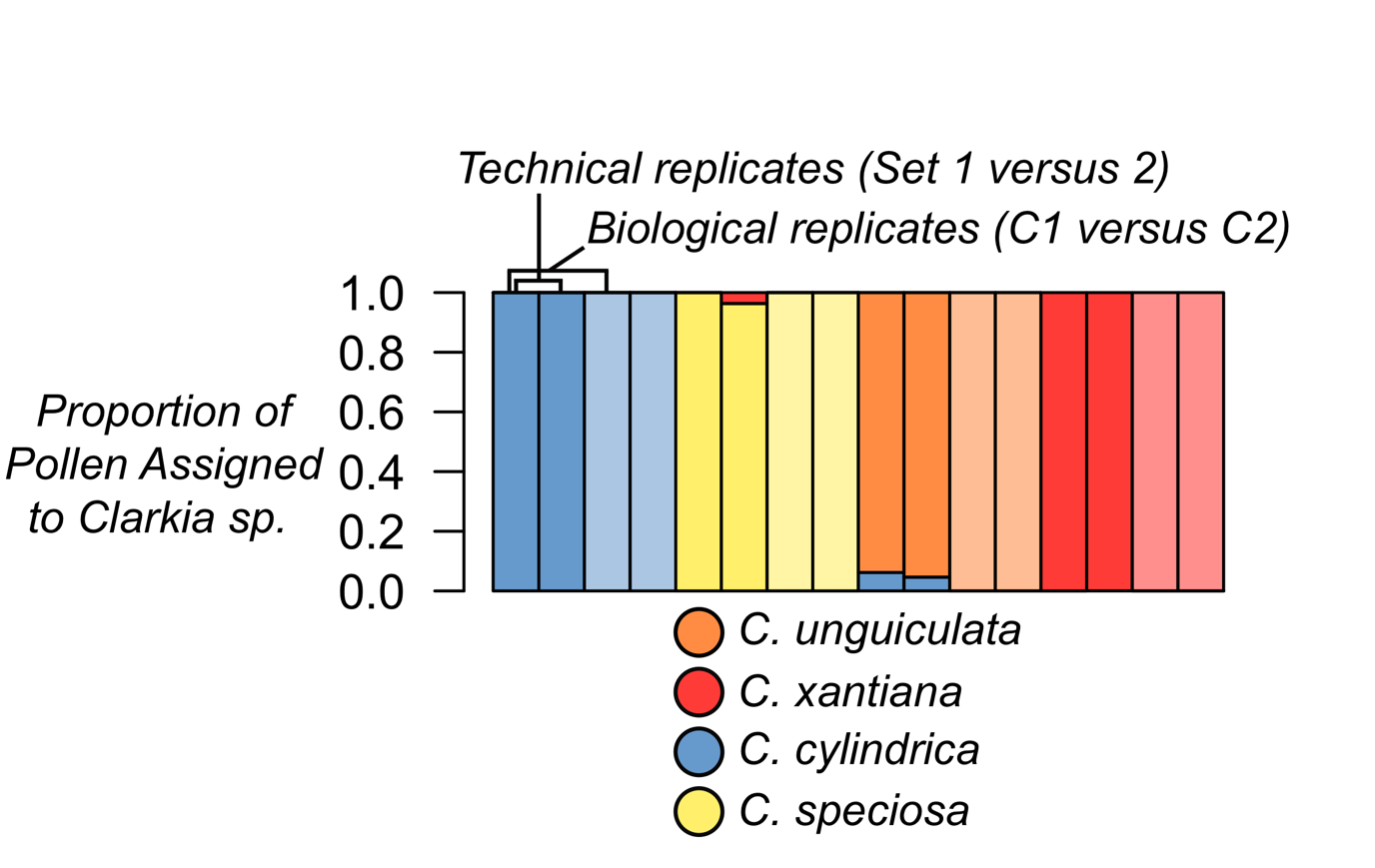

**Figure S1.** Proportion of pollen assigned to each of the four species of *Clarkia* sampled from pollen of known origin (i.e. unlikely to have a mix of >1 species). We ran two known samples (biological replicates) from each of the four species across two sets of samples (*i.e.* technical replicates between sets).

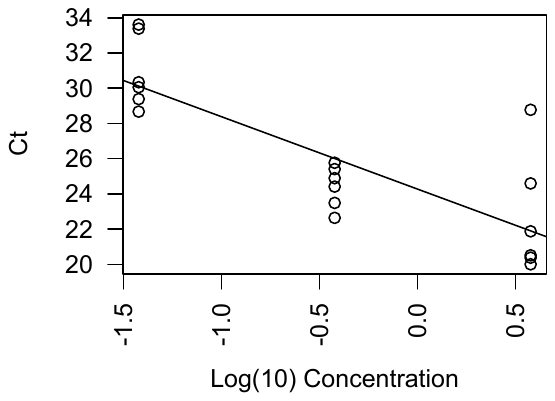

**Figure S2.** Standard curve of dilutions (1:10, 1:100, and 1:1000) of known sample DNA concentration (total DNA concentration of extract = 37.8 ng/uL) and calculated C_T_. The linear regression shows the relationship used to transform C_T_ values from unknown DNA samples into DNA concentration estimates: [log_10_(DNA)] = (C_T_-24.273)/(-4.1112). The R^2^ = 0.65 suggests there is moderate variance among the samples, which could be due to pipetting error, particularly for the highest concentration samples (e.g. log_10_[DNA] = 0.58). Removing one replicate in this case increases the R^2^ to 0.82. However, this variation should not change the overall relative abundances among samples.

*A note on TaqMAN Probe Sensitivity*

Our method of quantitative amplicon sequencing (below) applies the general theory of quantitative PCR. Before introducing this method, a natural question regarding our approach is why we did not use fluorescence-based quantitative PCR, such as TaqMan probes (Thermo Fischer Scientific, Waltham MA, USA). We note here that we did develop and test TaqMan probes for the small region that distinguished the four *Clarkia* species within the manufacturers recommended design specifications. However, these probes, with 1-4 species-specific SNPs, were not sensitive enough to fluoresce exclusively enough in the target species, and therefore did not allow us to distinguish among any of the four *Clarkia* species. Thus, the TaqMan chemistry was not sensitive enough to generate reliable relative abundance information. We are unaware of other published reports discussing this sensitivity, which was also not clear to the manufacturer, and thus raise this point here for researchers interested in applying fluorescent probes to quantify relative abundance using a small number of SNPs.
